# Critical minimum temperature limits xylogenesis and maintains treelines on the Tibetan Plateau

**DOI:** 10.1101/093781

**Authors:** Xiaoxia Li, Eryuan Liang, Jozica Gricar, Sergio Rossi, Katarina Cufar, Aaron M. Ellison

## Abstract

Physiological and ecological mechanisms that define treelines are still debated. It is suggested that the absence of trees above the treeline is caused by the low temperature that limits growth. Thus, we raise the hypothesis that there is a critical minimum temperature (CTmin) preventing xylogenesis at treeline. We tested this hypothesis by examining weekly xylogenesis across three and four growing seasons in two natural Smith fir (*Abies georgei* var. *smithii*) treeline sites on the south-eastern Tibetan Plateau. Despite differences in the timing of cell differentiation among years, minimum air temperature was the dominant climatic variable associated with xylem growth; the critical minimum temperature (CT_min_) for the onset and end of xylogenesis occurred at 0.7±0.4 °C. A process-based-modeled chronology of tree-ring formation using this CT_min_ was consistent with actual tree-ring data. This extremely low CT_min_ permits Smith fir growing at treeline to complete annual xylem production and maturation and provides both support and a mechanism for treeline formation.

## INTRODUCTION

The explanations for treeline formation focus on limitations of available resources (Stevens & Fox 1991; Susiluoto *et al.* 2010), establishment sites (e.g., Smith *et al.* 2003), or time available for growth (e.g., Körner 1998), although this ecophysiological causes still remain debated (Tranquillini 1979; Holtmeier 2003; Körner 2003; Cui *et al.* 2005). Based on notable similarities in temperatures at treelines (Körner & Paulsen 2004), Körner (1998) proposed that low temperatures limit the time available for meristematic growth and cell division (Körner 1998). This hypothesis has been supported by phenomenological data. For example, treeline trees tend to have higher amounts of non-structural carbohydrates than trees growing at lower elevation, suggesting that treeline are limited more by growth processes than by photosynthesis and carbon assimilation (e.g. Hoch & Körner 2003; Shi *et al.* 2008; Fajardo *et al.* 2012). In parallel, dendroclimatic studies have identified a signal of reduced growth during periods with low temperatures at treelines in cold and humid areas (e.g., Mäkinen *et al.* 2000; Oberhuber 2004; Frank & Esper 2005; Ettinger *et al.* 2011).

Physiological manifestations of the growth limitation hypothesis include a constraint on the production of new cells by meristems below a critical minimum temperature (CT_min_) (Körner 1998) and a trade-off between taking maximal advantage of the length of the growing season while avoiding cellular damage due to early (fall, winter) or late (winter, spring) freezing events (Chuine & Beaubien 2001; Kollas *et al.* 2014). Such a trade-off would suggest a narrow thermal window for the onset and cessation of xylem formation at treeline. Indeed, recent studies focused on temporal dynamics of xylem formation in various tree species at treeline (Rossi *et al.* 2007; Seo *et al.* 2008; Gruber *et al.* 2009; Moser *et al.* 2010; Lenz *et al.* 2012). One study reported that a gradual increase in temperature (heat sum) was associated with the onset of cambial activity (Seo *et al.* 2008), whereas another estimated a CT_min_ of 5.6–8.5°C for xylogenesis at the altitudinal treeline in the Eastern Alps (Rossi *et al.* 2007). Separating gradual (heat-sum) and threshold (CT_min_) effects on xylogenesis at treeline has not yet been accomplished.

A mechanistic model can provide a deeper understanding on a climatic control on tree growth dynamics. The process-based forward model, such as, Vaganov-Shashkin (VS) model, was used to simulate the climatic control on conifer tree-ring growth (Vaganov et al. 1999; Anchukaitis et al. 2006; Evans et al. 2006). The critical temperature for cambial activity is a key parameter to model tree growth despite that its observed data are less available as a model input.

Our observations at the upper Smith fir *(Abies georgei* var. *smithii)* treeline of the south-eastern Tibetan Plateau, including a decade of uninterrupted *in situ* micrometeorological measurements and weekly collection of microcores containing cambium and forming wood during three consecutive growing seasons provides an opportunity to examine both gradual and threshold effects of temperature on xylogenesis at a natural alpine treeline. Specifically, we tested the potential for thermal control of xylogenesis to be a mechanism underlying the growth limitation hypothesis by (1) identifying the timing and dynamics of xylem formation in Smith fir growing at treeline as a function of climatic factors; and (2) detecting a plausible CT_min_ for xylogenesis. Previous studies have found that its growth near treeline is constrained by the minimum temperature in summer (Bräuning & Mantwill 2004; Liang *et al.* 2009). The onset of bud swelling and needle unfolding in Smith fir is delayed by 3.5 days per 100 m increase in elevation (Wang *et al.* 2013), indicating a thermal limitation of tree phenology. Therefore, we hypothesized that minimum temperature limits xylem formation and that a threshold minimum temperature controls the timing of the onset and ending of xylem formation. If this potential CT_min_ is reasonable, as a primary temperature parameter, it will be expected to be used in Vaganov-Shashkin (VS) model of tree-ring formation to simulate regional tree growth at Smith fir treeline on the south-eastern Tibetan Plateau.

## MATERIALS AND METHODS

### Study sites and tree selection

The study focused on the natural alpine treeline of Smith fir *(Abies georgei* var. *smithii)* growing on the eastern side of the Sygera Mountains (29° 10′ − 30° 15′ N, 93° 12′ – 95° 35′ E) on the south-eastern Tibetan Plateau (Liang *et al.* 2011). The south-eastern Tibetan Plateau is characterized by a cold and humid climate, and has the highest natural treeline (up to 4900 m a.s.l.) in the Northern Hemisphere (Miehe *et al.* 2007). Smith fir *(Abies georgei* var. *smithii)* is one of the dominant treeline species in this region. This tree is a shade-requiring conifer and the upper treeline depends on topographic aspect and ranges from 4,250 to 4,400 m a.s.l. We studied two sites at open-canopy treelines: Site 1, was at 4360 m a.s.l. on an east-facing slope, and Site 2, was at 4250 m a.s.l. on a south-east-facing slope. The sites were 200 m apart, on slopes < 15° *Rhododendron aganniphum* var. *schizopeplum* dominated the understory. The coverage of Smith fir was < 20% and the podzolic soils had an average pH value of 4.5.

At each site, five dominant trees were selected in April 2007. These trees had a mean age of 201 ± 24 and 117 ± 14 years, and mean diameters at 1.3-m above ground of 34 ± 4 and 44 ± 7 cm in Sites 1 and 2, respectively. Because repeated sampling could cause severe wounding that could modify xylem formation, another five trees per site with homogeneous diameters were chosen for the samplings in 2009 and 2010. Trees with polycormic stems, partially dead crowns, reaction wood, or other evident damage were avoided.

### Meteorological data

An automatic weather station (Campbell Scientific, CR1000) was installed in November 2006 in an open area above the treeline (29°39′ N, 94°42′E, 4390 m a.s.l.) at a linear distance of ≈150 m and 200 m from Sites 1 and 2, respectively. Measurements of air (3-m above ground) and soil temperature (at 10-, 20- and 40-cm depths), precipitation, snow fall, and soil water content (at 10-, 20-, and 40-cm depths) were collected at 30-minute intervals. These data were used to compute daily averages, minima, and maxima of each variable.

### Microcoring and histological analyses

Xylem formation was studied from 2007 until 2010 at Site 1 and from 2007 to 2009 at Site 2. One microcore (15-mm long, 2-mm diameter) was collected from each tree weekly from May until October around the stem at breast height using a Trephor tool. Immediately after removal from the trees, the microcores were fixed in a formalin-ethanol-acetic acid (FAA) solution. The microcores contained innermost phloem, cambium, developing xylem, and at least three previous xylem growth rings. In the laboratory, the microcores were dehydrated with successive immersions in a graded series of ethanol and *d*-limonene, then embedded in paraffin. Transverse sections (9-12 μm in thickness) were cut from the samples with a Leica RM 2245 rotary microtome using Feather N35H knives (Osaka, Japan). Sections were stained with a mixture of safranine (0.5 % in 95 % ethanol) and astra blue (0.5 % in 95% ethanol) and observed with a Nikon Eclipse 800 light microscope under bright field and polarized light to identify the phases of differentiation of the developing xylem cells. In cross-section, cambial cells were characterized by thin cell walls and small radial diameters (Deslauriers *et al.* 2003). Newly-formed xylem cells in the phase of cell enlargement contained protoplasts, had thin primary walls, and a radial diameter at least twice the size of the cambial cells. The onset of cell-wall thickening was determined by birefringence in the cell walls under polarized light. Mature cells had completely red-stained walls and empty lumen. For each sample, the total current xylem cell number was determined by counting the number of cells undergoing enlargement, cell-wall thickening, and the number of mature cells along three radial files (Deslauriers *et al.* 2003).

### Data standardization and estimating the rate of xylem formation

The data were standardized to compensate the variation in the number of xylem cells along the tree circumference. The total cell number of the previous years was counted on three radial files per sample and used for standardization. The standardized number of cells *ncij* number in the *i*^th^ phase of the *j*^th^ sample was calculated as:

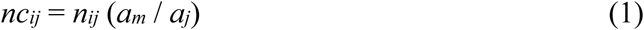

where *nij* is the number of cells in the current year, *a_m_* is the mean number of cells of the previous ring of all *j*-samples and a_*j*_ is the mean number of cells of the previous ring in each *j*-sample.

We modelled the dynamics of xylem formation by fitting a Gompertz function to the number of xylem cells that were produced through time:

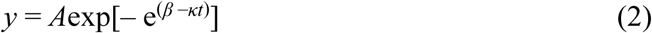

where *y* is the weekly cumulative sum of tracheids, *t* is the time of the year computed as day of the year, *A* is an asymptote (constant) and *β* and *k* are constants reflecting the *x*-intercept placement and rate of change, respectively (Deslauriers *et al.* 2003). Model parameters were estimated using the Origon software package (Version 8.5, OriginLab Corporation, Northampton, MA, USA).

### Estimation of the onset and ending of xylem formation

We used directly observations of cell differentiation to identify the onset, ending, and duration of xylem formation from counts of the number of cells in three radial files per tree. In spring, xylem formation was considered to have started when at least one tangential row of cells was observed in the enlarging phase. Because of the weekly resolution of the monitoring, we used the occurrence of 1-2 enlarging xylem cells along any of the checked three radial files as an indicator the xylem formation had begun (Li *et al.* 2013). In late summer, when cells were no longer observed in the wall thickening and lignification phase, xylem formation was considered to have ended. The duration of xylogenesis was estimated as the number of days between the dates of onset and ending of xylem formation; we report the average among trees for each studied site and year.

Comparisons between sites in onset, duration, and ending of differentiation in the developing xylem ring, were done with generalized linear models (GLM). Homoscedasticity was checked using Shapiro-Wilk and Levene tests.

### Identifying CT_min_

Logistic regression (LOGISTIC procedure in SPSS 16.0) was used to model the probability of xylogenesis as a function of air temperature. Xylem cell production was coded zero (not occurring) or one (occurring). CT_min_ was estimated as that temperature when the probability of ongoing xylem growth equalled 0.5 (Rossi et al. 2008). CT_min_s also were calculated for 5% and 95% of the maximum number of xylem cells. For each tree and year, the model was fitted with three respective daily temperature series (mean, absolute minimum, and absolute maximum). Model verification included the likelihood-ratio *χ*^2^, Wald’s *χ*^2^ for regression parameter and goodness of fit, and Hosmer-Lemeshow *Ĉ* for possible lack of fit. None of the models were excluded because of a lack of fit. CT_min_s were compared between sites and years using analysis of variance (ANOVA) models. Model validation was performed by comparing the observations with the predicted values when using estimated CT_min_s.

### Climate-growth relationships

We used two approaches to identify relationships between intra-annual xylem growth and climatic variables during four growing seasons. One approach consisted of computation of Pearson’s correlation coefficients between xylem cell production and weather data for weekly intervals. Weather data here include daily mean, daily absolute minimum, daily absolute maximum temperatures, growing degree-days (GDD) > 5°C, and sums of precipitation.

Intra-annual xylem growth may be controlled both by endogenous (e.g. hormonal regulation) and exogenous factors (e.g., climate). To analyse the climatic effect, a common approach was used to remove the endogenous growth trend by fitting a growth curve, and to estimate the growth departure, calculated as the dimensionless ratio between observed and expected growth (Fritts 1976). This ratio (hereafter called ‘growth index’) was calculated as the number of tracheids produced during the week divided by the expected values estimated using the Gompertz function (Zhai *et al.* 2012). To account for possible effects of time-lags, daily weather data were averaged (temperature) or summed (precipitation) weekly from 1 to 10 d prior to each sampling date (referred to as P1 to P10). To minimize the effects of temporal autocorrelation, correlation coefficients were calculated on first-order differences for both datasets.

### Tree-ring modeling

We used the Vaganov-Shashkin (VS) model to simulate tree-ring growth at the Smith fir treelines in the Sygera Mountains. The VS model estimates xylem formation and its internal characteristics based on equations relating daily temperature, precipitation, and sunlight to the kinetics of xylem development (Vaganov *et al.* 2006). It assumes that climatic influences are directly but nonlinearly related to tree-ring characteristics through controls on the rates of cambial activity processes. To date, it has been successfully used to simulate and evaluate the relationships between climate and tree-ring formation under a variety of environmental conditions in many different regions (Anchukaitis *et al.* 2006; Evans *et al.* 2006; George *et al.* 2008; Shi *et al.* 2008; Zhang *et al.* 2011; Gou *et al.* 2013). Variables used as input for the VS model included soil moisture, depth of root system, temperature sum for initiation of growth, soil water drainage rate, and maximum daily precipitation falling into soil were taken from field observations. For the parameter of minimum temperature, we used the abovementioned CT_min_. Model fit was evaluated against an actual high-quality tree-ring width chronology from Smith fir treeline in the Sygera Mts., which was developed and used for paleoclimatic reconstructions in this region (Liang *et al.* 2009). The best estimate of physiological CT_min_ was found by iteration and comparison between simulated and observed chronologies (1960 – 2006).

Finally, a single simulated tree-ring width chronology was created for the Smith fir treeline in the Sygera Mts. based on daily climate data from Nyingchi meteorological stations (3,000 m a.s.l.). To account for the altitude differences between Nyingchi and the study sites, we extended the time series of daily temperatures at the treeline back to 1960 based on a linear regression of the Nyingchi data and our own micrometeorological data (r^2^ ≥ 0.89, 2007-2010).

## RESULTS

### Micrometeorological conditions at the upper treeline

The sampling sites at the upper treeline were cold and humid. In spite of a difference of 110 m in elevation and different topographical aspects of the two treeline sites, they had fairly similar temperatures (Supporting Information Fig. S1). Annual temperatures (2007 – 2010) ranged from 0.1 to 0.9 °C, while growing-season (June-September) temperatures ranged from 6.4 to 7.1 °C (Fig. 1). On average, the annual precipitation was 951 mm, of which 62 % fell during the monsoon season (June to September). Snowfall mainly occurred from November to May. Due to snowmelt and increased precipitation, soil moisture content increased rapidly from the beginning of April and remained above 30 % from early May until November, and finally decreased to near 0 in late November and early December. The year 2008 was characterized by heavy spring snowfall and had the latest snowmelt and soil thawing during the four studied years (Fig. 1).

**Figure 1.**
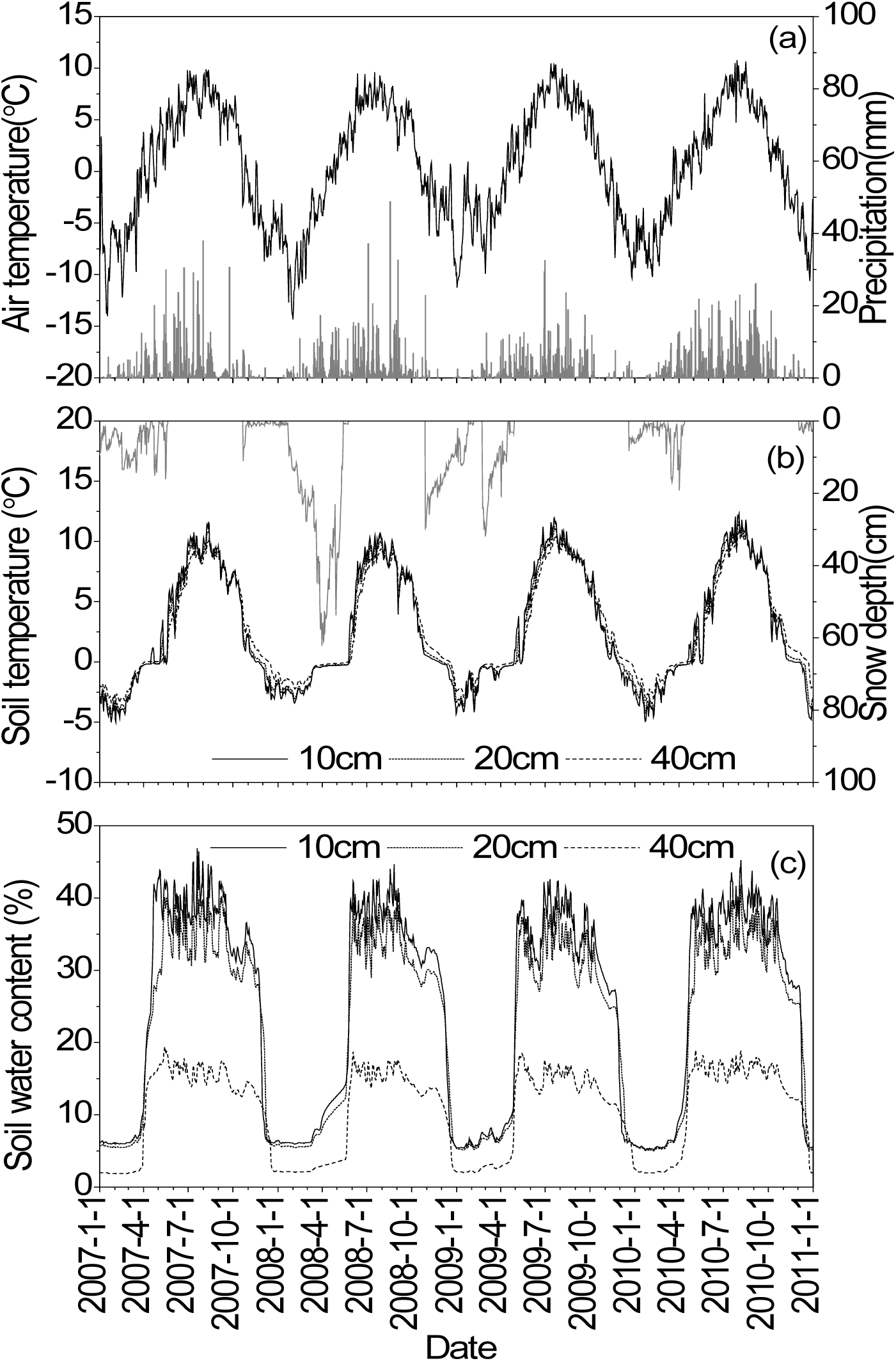
Micrometeorological conditions (2007 – 2010) at the upper treeline in the Sygera Mts., SE Tibetan Plateau, showing (a) daily mean air temperature and daily total precipitation, (b) daily soil temperature (at depths of 10, 20 and 40 cm) and snow depth, and (c) daily mean soil volumetric moisture contents (at depths of 10, 20 and 40 cm).

### Xylem formation

The onset of xylem formation occurred from late May to early June and differed significantly among years (*F* = 15.73, *P* < 0.001). The onset of xylem formation was observed 4 - 9 days later in 2008 than in the other years, at both sites (Fig. 2 a). No difference was found in onset of xylogenesis between sites (*F* = 2.31, *P* > 0.05). Xylem formation ended between the beginning and the end of September and differed significantly among years (*F* = 10.42, *P* < 0.005), and 1-2 weeks later in 2010 at site 1 (Fig. 2 b).

**Figure 2.**
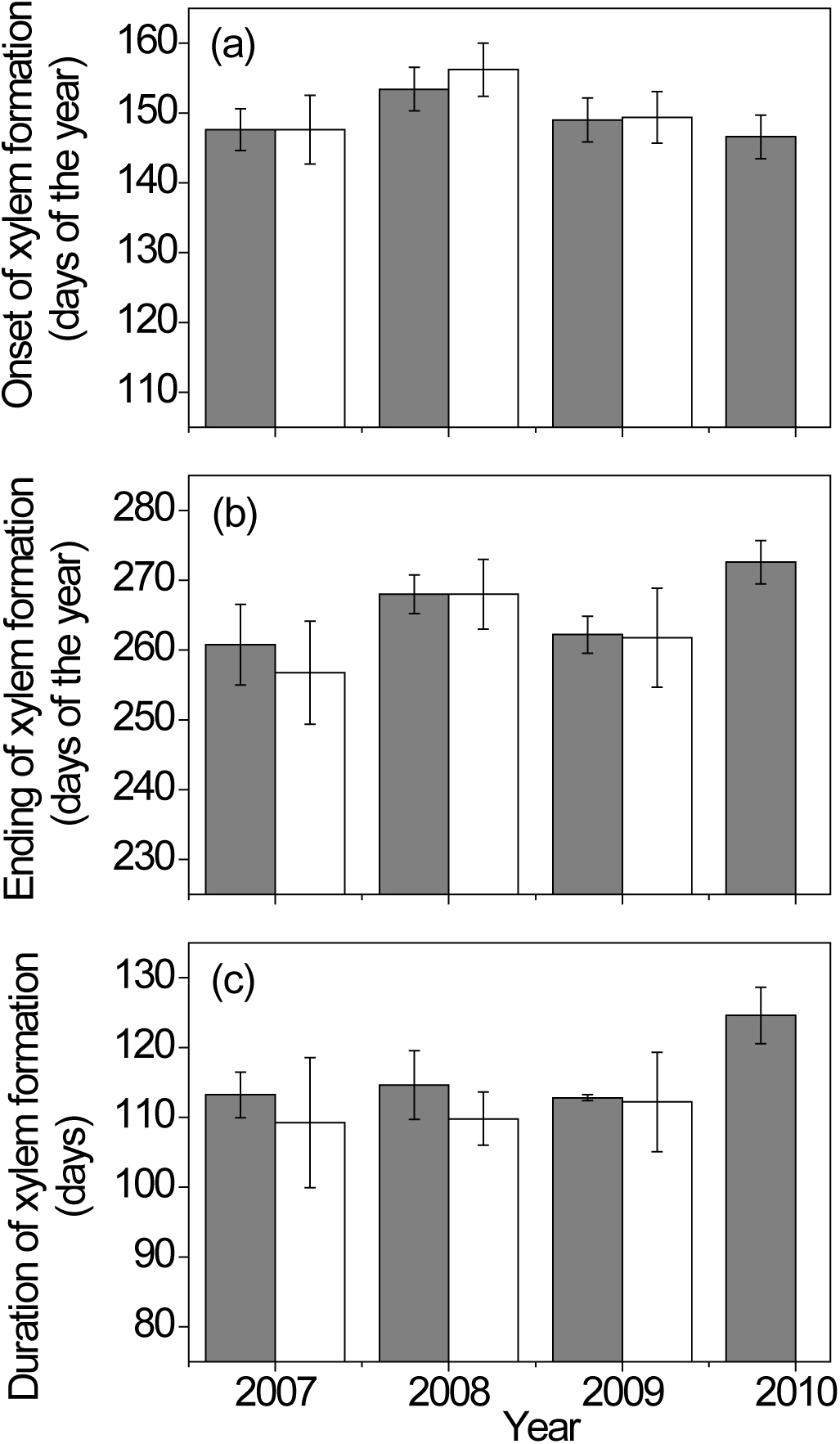
Onset (a), ending (b) and duration (c) of xylem formation of Smith fir (*Abies georgei* var. *smithii*) based on weekly xylogenesis observations at Site 1 (4360 m a.s.l.) (grey columns) and Site 2 (4250 m a.s.l.) (white columns). Error bars indicate standard deviation among trees.

Overall, the duration of xylem formation lasted from 109 to 125 days (Fig. 2 c), with no significant differences detected between sites (*F* = 3.80, *P* > 0.05). Conversely, there were significant variations among years (*F* = 4.71, *P* < 0.05). From 2007 to 2009, the average period between the onset and ending of xylem formation was 107 days, whereas an average of 118 days was required to complete xylem formation in 2010.

### Relationship between climate and xylem formation

Weekly cumulative xylem production was fit well by the Gompertz function (0.96 ≤ r^2^ ≤ 0.98; Supporting Information Table S1 and Fig. S2). Intra-annual xylem cell production was significantly and positively correlated with daily minimum and mean air temperatures and GDD > 5 °C at both sites (Fig. 3 a, b). However, only minimum temperature was significantly correlated with growth indices after removing the growth trends (Fig.3 c, d). At site 1, positive correlations between growth indices and minimum temperature were found for time lags of 0 – 3 days (*r* = 0.34, *P* <0.05), whereas the corresponding time lags were 7 – 10 days at site 2 (*r* = 0.42, *P* <0.05). No significant correlations were found between xylem cell production or growth index and precipitation from P0 to P10.

**Figure 3.**
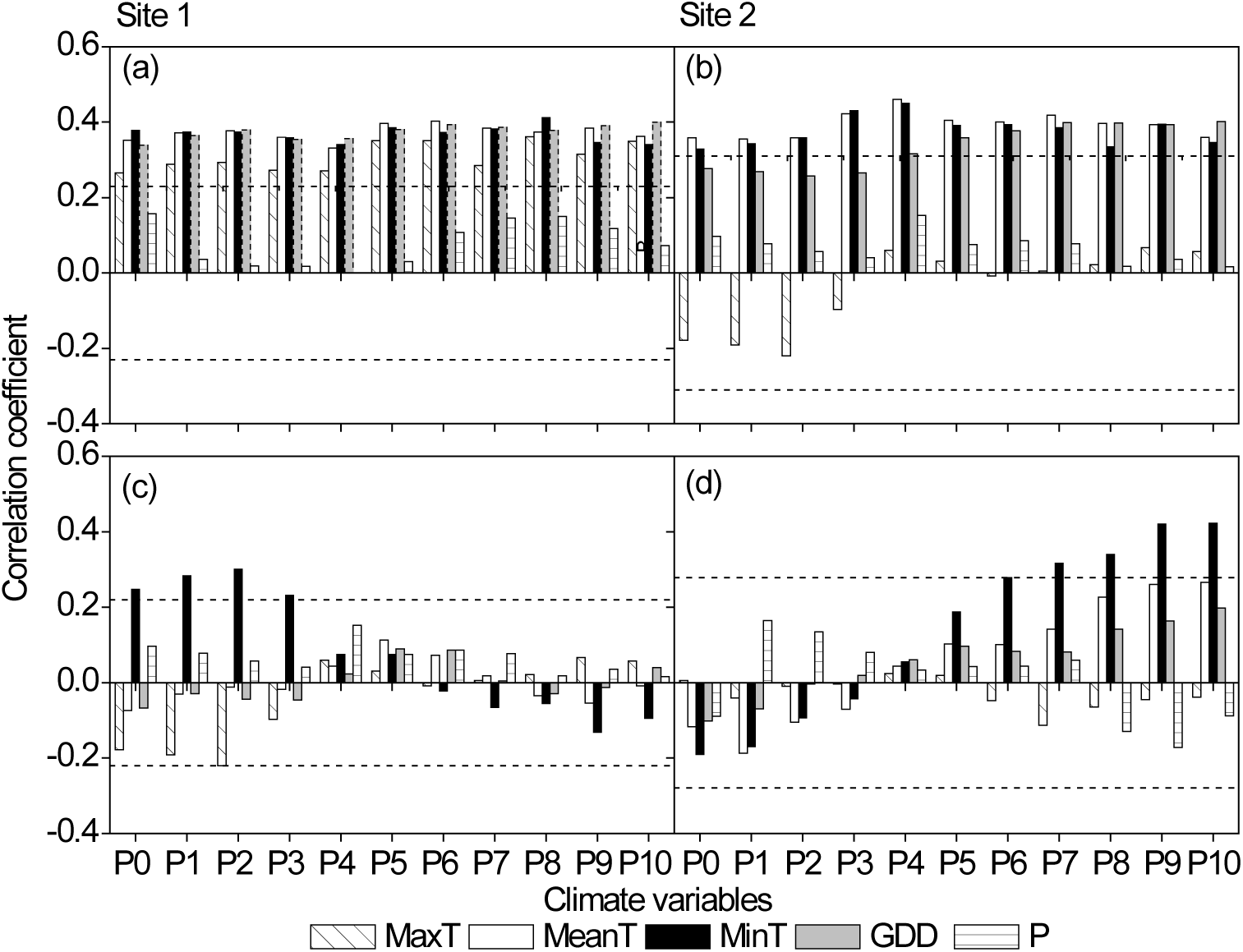
Lagged (0 - 10 days) Pearson correlation coefficients between xylem cell production (a, b), growth index (c, d) and corresponding climatic variables. P0 represents the weekly climatic mean for the exact period between two sampling dates. P1 to P10 represent the weekly means lagged 1 - 10 days before the sampling date. Dotted horizontal lines show the 95% confidence limits. *Abbreviations*: MaxT = maximum temperature, MeanT= mean temperature, MinT= minimum temperature, P = precipitation and GGD = growing degree days above 5°C.

### CT_min_

The critical minimum air temperature (CT_min_) at which there was a 0.5 probability that xylem formation was ongoing is shown in Fig. 4 and Table 1 for Site 1 (2007 – 2010) and Site 2 (2007 – 2009). Based on direct observations of xylogenesis, the values respectively for minimum, mean, and maximum temperatures of 0.6, 4.0, and 9.3°C were estimated for the onset of xylogenesis, while the corresponding values for the ending of xylem differentiation were 0.7, 3.9, and 9.0 °C. There were no differences among critical temperatures for the onset and ending of xylogenesis (ANOVA, *P*>0.05), with values of 0.7 ± 0.4, 3.9 ± 0.5, and 9.1 ± 0.6 °C for the minimum, mean and maximum temperatures, respectively. No significant differences were found between the two sites in terms of the estimated air temperature thresholds for the onset and ending of xylogenesis (ANOVA, *P* > 0.05). The mean air temperature during the period of xylem formation based on weekly observations at both sites was 6.8 ± 0.4°C.

**Figure 4.**
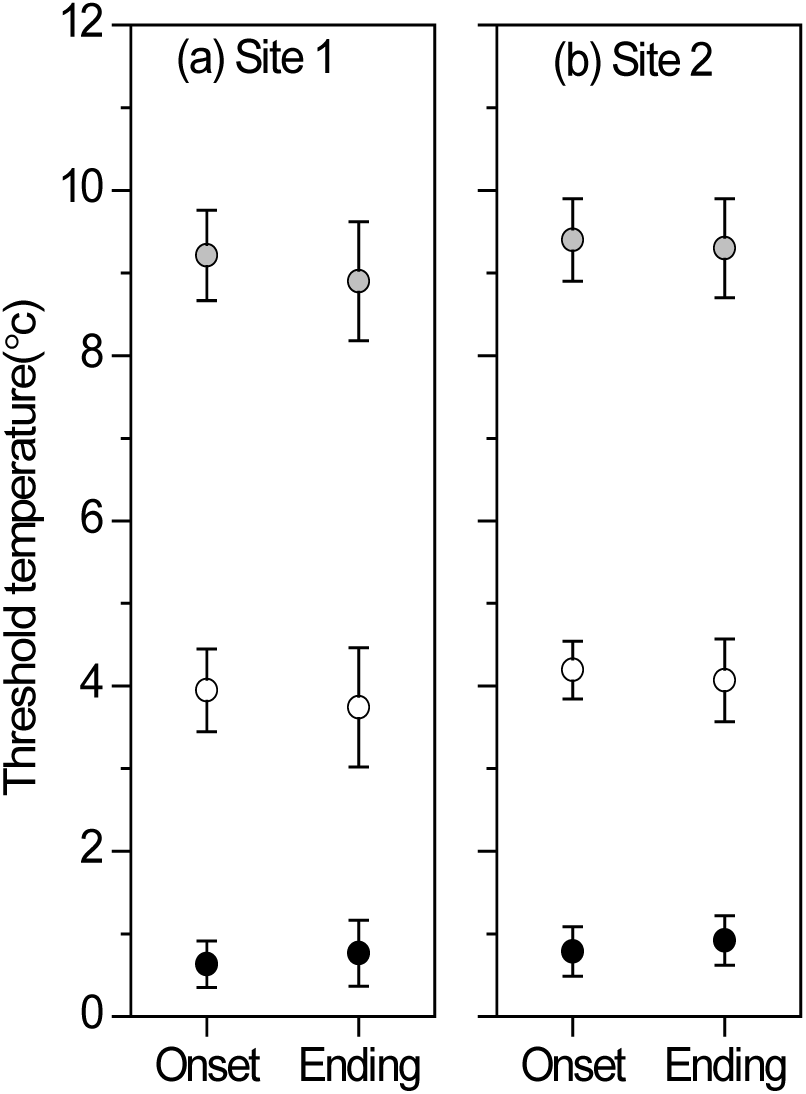
Critical minimum (black dots), mean (white dots) and maximum (grey dots) air temperatures at Sites 1 and 2, corresponding with the 0.5-probability of the onset and ending of xylem formation according to xylogenesis observations in Smith fir. Error bars indicate the standard deviation among trees.

**Table 1.**
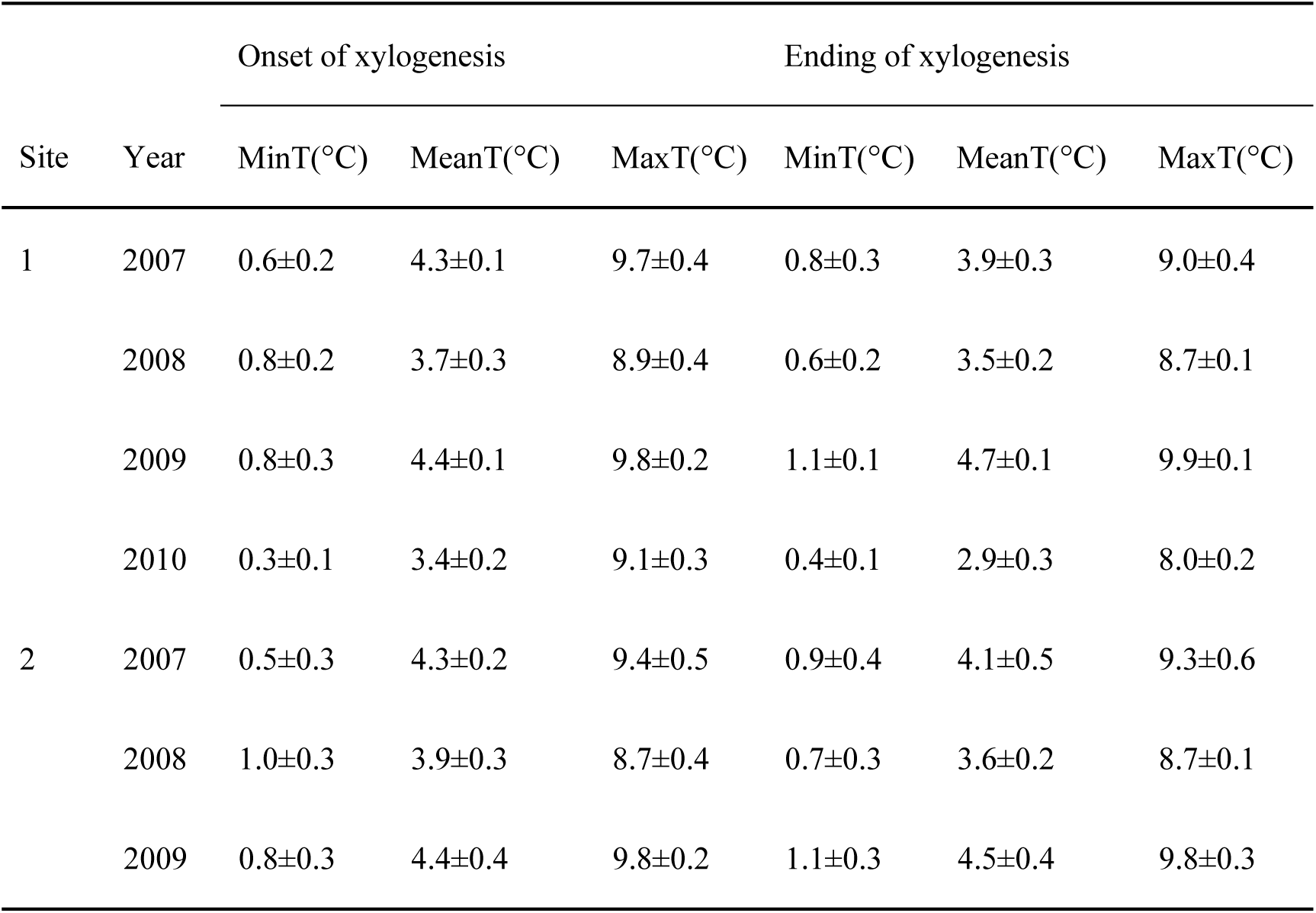
Mean ± standard deviation of the threshold daily maximum (MaxT), mean (MeanT) and minimum (MinT) temperatures for the onset and ending of xylogenesis.

### Tree-ring modeling

Applying the Vaganov-Shashkin (VS) model starting with an estimated CT_min_ = 0.7 °C, the best estimate of physiological CT_min_ was 0.9 °C (Table 2, Fig. 5). The correlation between observations and predictions varied slightly at CT_min_ of 0.3-0.9°C, while it descreased rapidly for values of CT_min_ >1°C (Fig. 6). Overall, significant, positive correlations were found between the modeled and real chronologies when CT_min_ varied within the range of 0.7 ± 0.4°C (*r* = 0.62, *P* < 0.01).

**Table 2.** The best estimate of parameters of VS model used in this study.

| Model parameter | Description(unit) | Value |
| --- | --- | --- |
| $CT_{min}$ | Minimum temperature for tree growth ($^{\circ}C$) | 0.9 |
| $T_{opt1}$ | Lower end of range of optimal temperatures ($^{\circ}C$) | 5.9 |
| $T_{opt2}$ | Upper end of range of optimal temperatures | 9.3 |
| $T_{max}$ | Maximum temperature for tree growth ($^{\circ}C$) | 19.9 |
| $W_{min}$ | Minimum soil moisture for tree growth (v/v) | 0.06 |
| $W_{opt1}$ | Lower end of range of optimal soil moisture (v/v) | 0.18 |
| $W_{opt2}$ | Upper end of range of optimal soil moisture (v/v) | 0.22 |
| $W_{max}$ | Maximum soil moisture for tree growth (v/v) | 0.50 |
| $T_{beg}$ | Temperature sum for initiation of growth ($^{\circ}C$) | 30 |
| $D_{root}$ | Depth of root system (mm) | 50 |
| $P_{max}$ | Maximum daily precipitation for saturated soil (mm) | 20 |
| $K_1$ | Fraction of precipitation penetrating soil (dimensionless) | 0.86 |
| $K_2$ | First coefficient for calculation of transpiration (mm/day) | 0.12 |
| $K_3$ | Second coefficient for calculation of transpiration ($1/^{\circ}C$) | 0.175 |
| $K_r$ | Coefficient for water infiltration from soil (dimensionless) | 0.006 |

**Figure 5.**
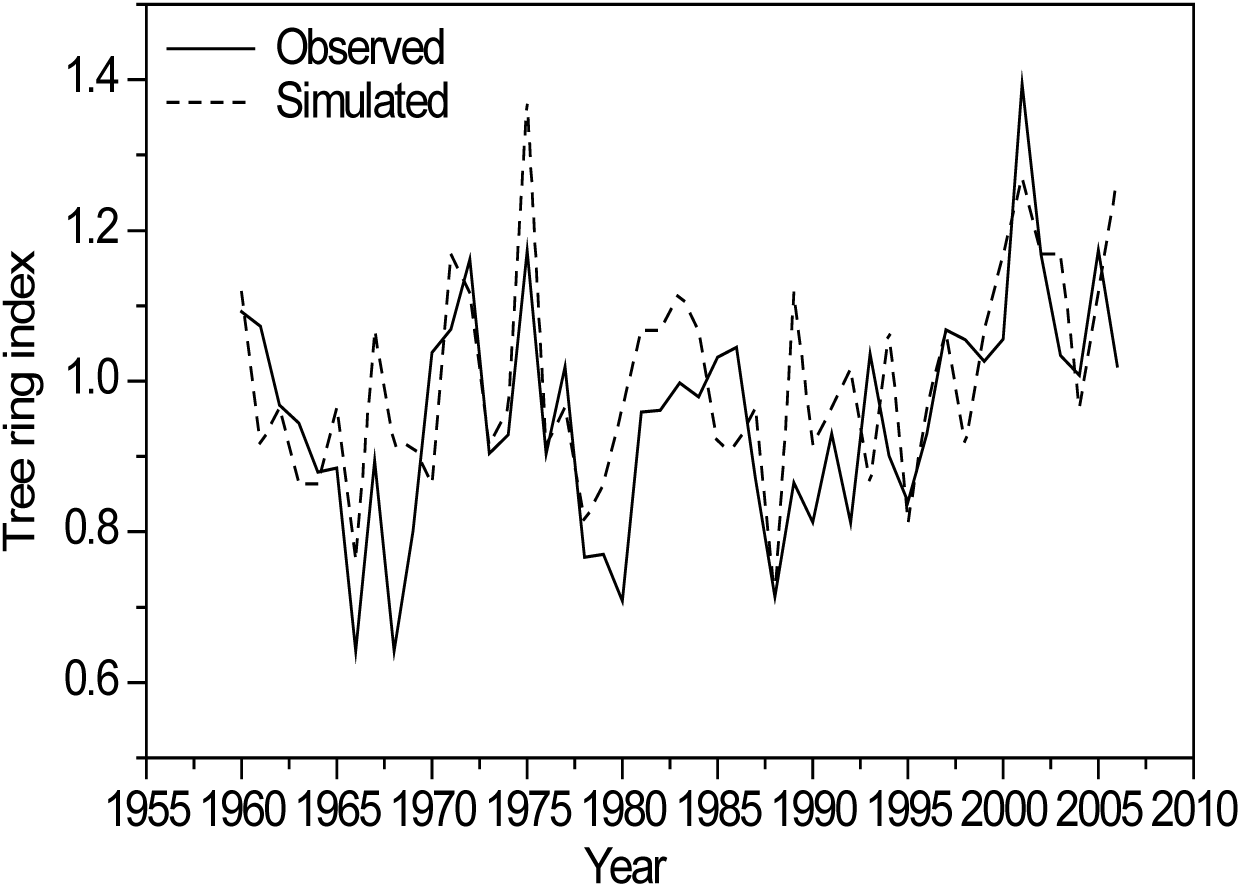
Observed (solid line) and simulated (dashed line) tree-ring width indices at Smith fir treeline in the Sygera Mts. on the south-eastern Tibetan Plateau, 1960 - 2006.

**Figure 6.**
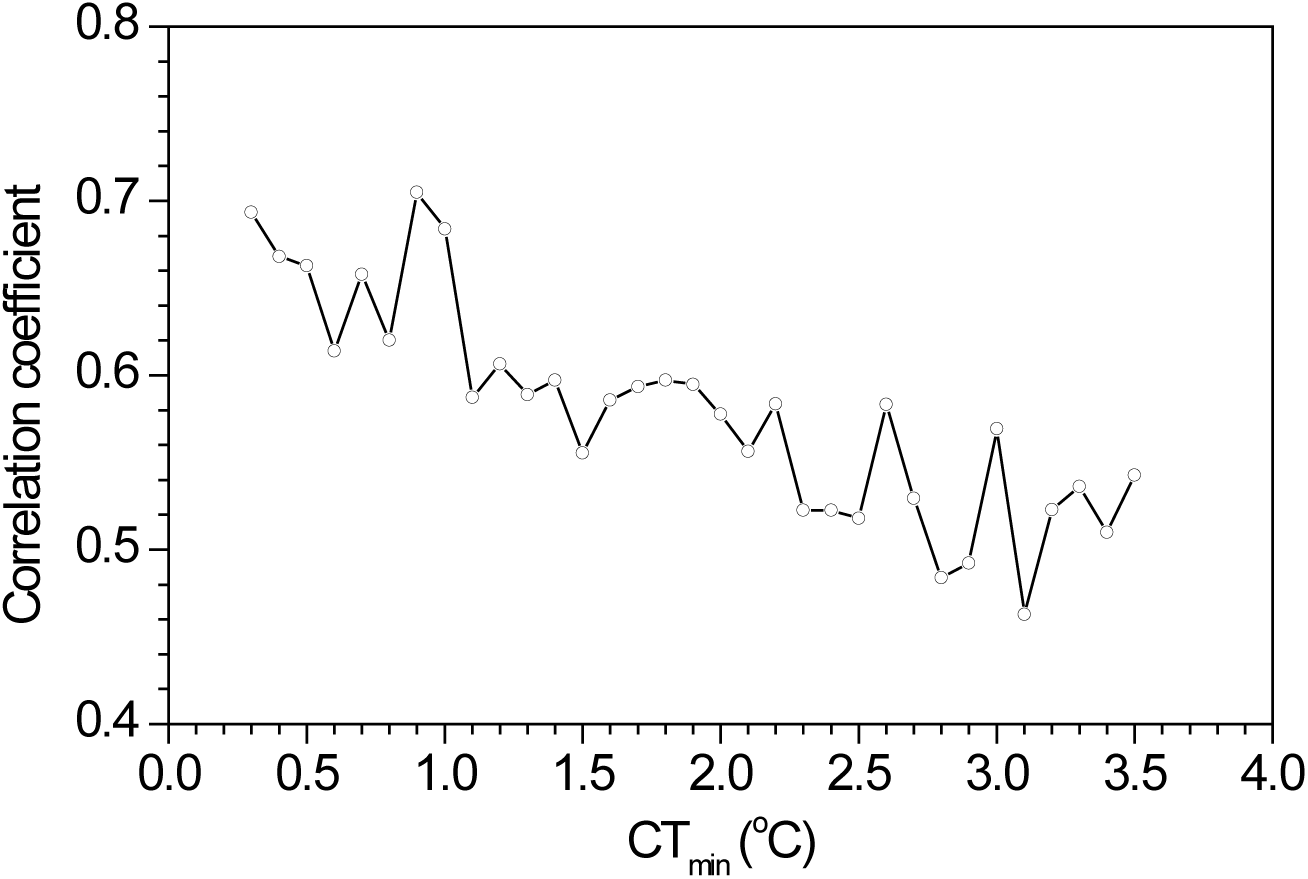
Pearson correlation coefficients between the observed and calculated values of tree ring width when the parameter of CT_min_ varied.

## DISCUSSION

The importance of temperature for xylem formation during and after its onset has been demonstrated repeatedly (e.g., Oribe *et al.* 2001; Gričar *et al.* 2006; Rossi *et al.* 2008). These and other data suggest that air temperature, not soil temperature, directly limits xylem formation at high latitudes and altitudes (Rossi *et al.* 2007; Lenz *et al.* 2012; Lupi *et al.* 2012). Minimum temperature is assumed to be an important driver of tree species range limits (Körner 2003; Kollas *et al.* 2014), and so a critical minimum temperature (CT_min_) with fairly narrow bounds should exist for the onset and ending of xylogenesis. However, long-term monitoring of xylem formation at natural treelines is limited, which has precluded assessment of CT_min_ for xylem formation by direct observations.

## Effects of climate on xylem formation

As predicted, minimum air temperature strongly limited xylem formation of Smith fir at the upper treeline on the south-eastern Tibetan Plateau. This finding accords with those from dendroclimatological analysis in the same study area (Liang *et al.* 2009) and xylem formation studies at high latitudes and altitudes (e.g., Rossi *et al.* 2008; Gruber *et al.* 2009). The importance of minimum air temperature may be related to the timing of cell differentiation, which perhaps occurs mainly during the night, when the temperature is lower (Hosoo *et al.* 2002, Steppe *et al.* 2015). Controlled experiments in Hamburg, Germany also showed that night temperatures could directly influence xylem cell expansion of *Podocarpus latifolius* (Dünisch 2010). According to Körner (2003), cell doubling time, which is highest and fairly constant at temperatures of 10-25°C, approaches infinity at 1-2°C, suggesting a minimum temperature limit on cell division. The simulated ring-width chronologies produced by VS model of tree-ring formation also exhibit similar positive correlations with the minimum temperature during summer (Supporting Information Fig. S3, *P*<0.01). CT_min_ is thus expected to limit xylogenesis of Smith fir at the treeline.

## Critical temperatures for xylem formation

Despite the variance in timing and duration of xylem formation during our four years of observations, minimum, average, and maximum temperatures for the onset and ending of xylogenesis were narrowly bounded with average values of 0.7, 3.9, and 9.1 °C, respectively. Most studies to date have indicated that xylogenesis in conifers growing in cold climates can take place when the daily minimum temperatures ≥ 4 – 5 °C (e.g. Rossi *et al.* 2008; Boulouf Lugo *et al.* 2012). However, based on the presented 4-year observations of xylogenesis and uninterrupted *in situ* micrometeorological measurements directly at the treeline, we found that the CT_min_ for xylogenesis in Smith fir is as low as 0.7 °C. In particular, based on this CT_min_, the modeled chronology produced by VS model is consistent with actual tree-ring data, suggesting that minimum temperature could be considered as a significant driver of xylem growth. Such a low CT_min_ may have evolved to provide sufficient time to complete xylogenesis at alpine treelines. The length of the growing season for stem growth diminishes with altitude and reaches a minimum at the alpine treeline. According to some authors (Rossi *et al.* 2008; Körner 2012), a tree can only survive when the length of growing seasons is at least 3 months and the mean growing season air temperature is 6.4.°C, since each of them critically constrains the growth and development of trees. At Smith fir treelines in south-eastern Tibet, the duration of xylem growth of 115 days provided by a CT_min_ < 1 °C, together with a mean air temperature of 6.8 °C during the growing season extended by this low CT_min_ meets these prerequisites for tree growth and development.

The dates of snow melting and soil thawing also are thought to be critical for the onset of xylogenesis and could therefore determine the annual xylem production (Vaganov *et al.* 1999). At our treeline sites, the onset of xylem growth occurred 7 - 30 days after soil thawing in spring, which coincided with the surpassing of CT_min_. This temporal lag might be related to a strategy of avoiding the risk of freezing damage during early cell development.

The growth limitation hypothesis predicts that the absence of trees above the treeline is attributable to critical minimum temperature for growth (Körner 1998). Treeline trees often have slower growth rates and higher non-structural carbohydrate levels than trees at lower altitudes (e.g. Hoch & Körner 2003; Shi *et al.* 2008; Fajardo *et al.* 2012), suggesting a carbon sink rather than carbon limitation. However, some authors have argued that tree populations with the highest non-structural carbohydrate concentrations may be the most carbon limited in terms of growth (Li *et al.* 2008; Wiley & Helloker 2012). Although our observations of xylogenesis cannot differentiate between carbon limitation and a carbon sink in Smith fir, the significant effect of a narrowly bounded CT_min_ on xylem cell production provides a physiological, rather than an ecological, mechanism for the growth limitation hypothesis.

## ACKNOWLEDGMENTS

This work was supported by the National Natural Science Foundation of China (41525001, 41130529, 41601204). International cooperation was supported by the bilateral project between China and Slovenia (BI-CN/09-11-012) and COST Action (FP1106, STReESS). AME’s participation in the project was supported by the Chinese Academy of Sciences President International Fellowship Initiative for Visiting Scientists, Grant no. 2016VBA074. We thank the Southeast Tibet Station for Alpine Environment Observation and Research, Chinese Academy of Sciences for the fieldwork and monitoring; Yongxiang Zhang from National Climate Centre, Beijing, China for her support on the VS model and Martin Cregeen for additional language editing.

## SUPPORTING INFORMATION

**Table S1.** Parameters of the Gompertz function (A, ß, K), *R*^2^ and day of the inflection point (tp) for Smith fir growing at two treeline sites, 2007 – 2010.

| | Year | $A$ | $\beta$ | $K(10^{-2})$ | tp | $R^2$ |
| --- | --- | --- | --- | --- | --- | --- |
| Site 1 | 2007 | 23.95 | 5.48 | 3.25 | 170.59 | 0.98 |
|  | 2008 | 22.97 | 5.97 | 3.32 | 179.32 | 0.96 |
|  | 2009 | 25.68 | 7.07 | 4.08 | 174.50 | 0.96 |
|  | 2010 | 21.08 | 5.67 | 3.25 | 176.67 | 0.99 |
| Site 2 | 2007 | 24.79 | 5.03 | 2.85 | 176.56 | 0.98 |
|  | 2008 | 21.22 | 6.97 | 3.87 | 178.13 | 0.98 |
|  | 2009 | 23.62 | 5.70 | 3.02 | 188.53 | 0.98 |

**Figure S1.**
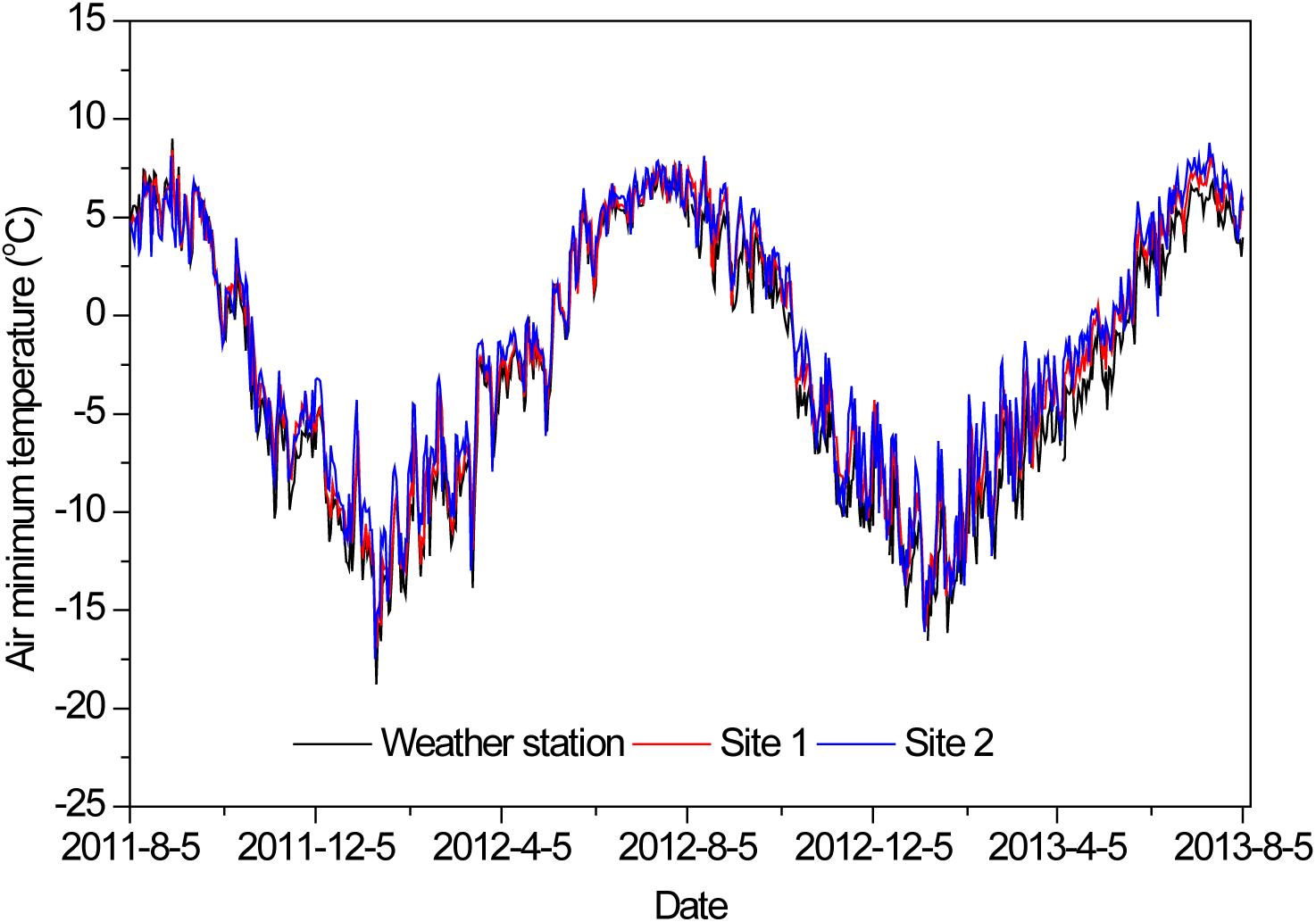
Minimum air temperatures recorded by the automatic weather station (black line) and temperature data logger at Site 1 (red line) and Site 2 (blue line) from August 5, 2011 to August 5, 2013.

**Figure S2.**
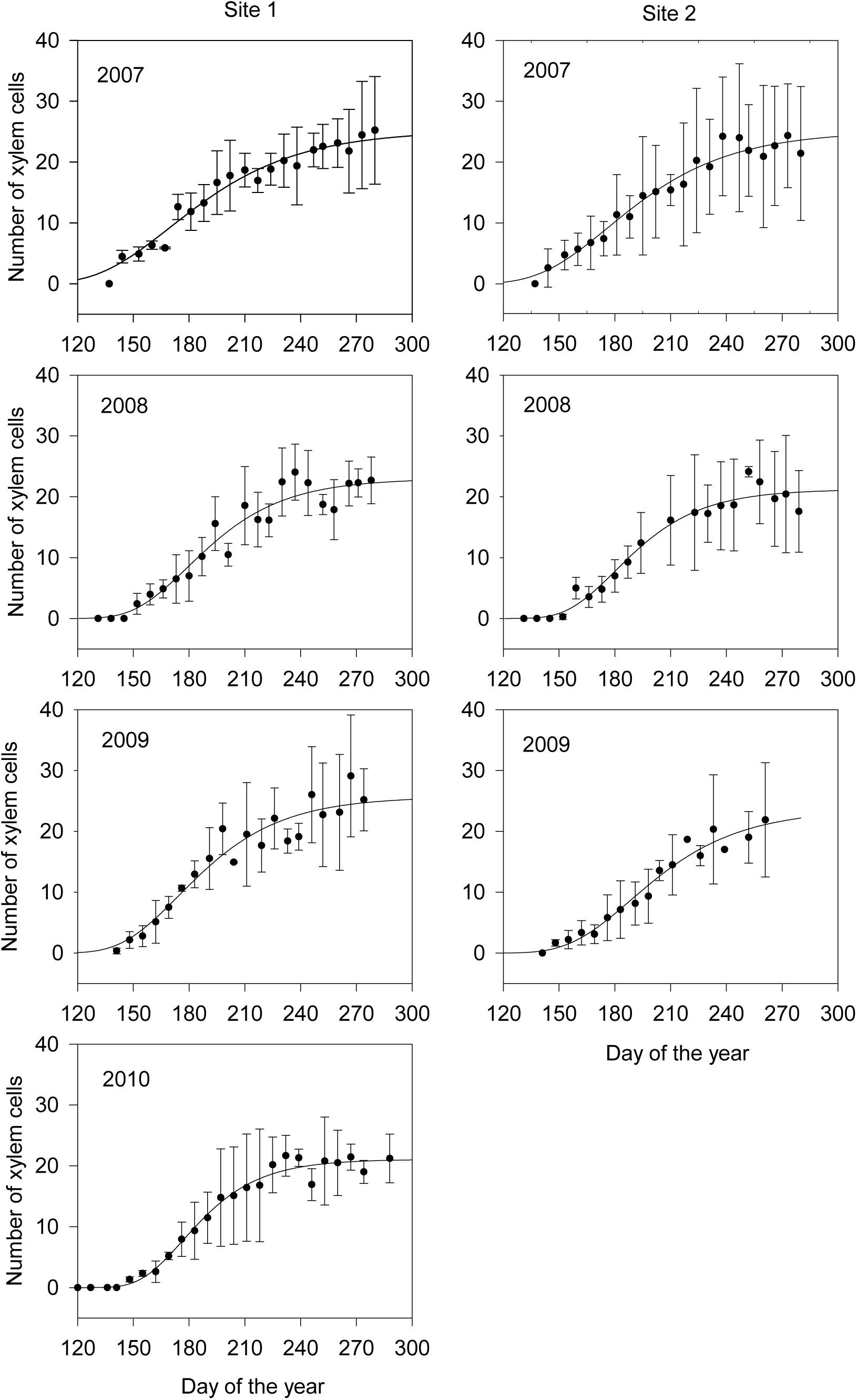
Dynamics of xylem growth (including enlarging, wall thickening and mature xylem cells) at two Smith fir treelines as modeled using a Gompertz function.

**Figure S3.**
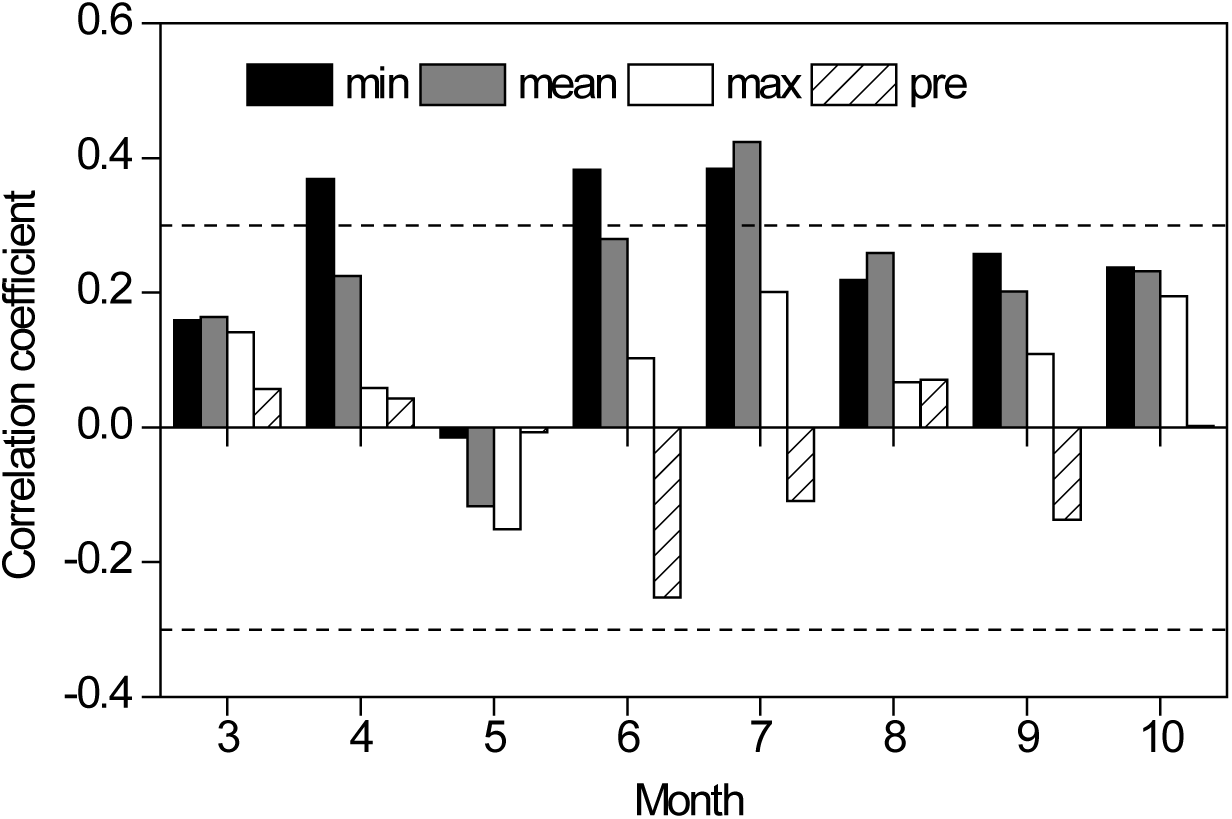
Correlations between the simulated tree ring chronology and monthly temperature and precipitation at Smith fir treeline in the Sygera Mountains on the south-eastern Tibetan Plateau. Dotted horizontal lines show the 95% confidence limits. *Abbreviations*: MaxT = maximum temperature, MeanT= mean temperature, MinT= minimum temperature and P = precipitation.

